# Regulation of Bacterial Two-Component Systems by Cardiolipin

**DOI:** 10.1101/2023.02.01.526740

**Authors:** Won-Sik Yeo, Sophie Dyzenhaus, Victor J Torres, Shaun R Brinsmade, Taeok Bae

**Affiliations:** Department of Microbiology and Immunology, Indiana University School of Medicine-Northwest, Gary, Indiana, 46408, USA; Department of Biology, Georgetown University, Washington, DC, USA; Department of Microbiology, New York University School of Medicine, New York, New York, USA

**Keywords:** *Staphylococcus aureus*, cardiolipin, SaeRS, two-component regulatory systems, SaeS sensor histidine kinase, virulence

## Abstract

The composition of phospholipid membranes is critical to regulating the activity of membrane proteins for cellular functions. Cardiolipin is a unique phospholipid present within the bacterial membrane and mitochondria of eukaryotes and plays a role in maintaining the function and stabilization of membrane proteins. Here, we report that, in the human pathogen Staphylococcus aureus, cardiolipin is required for full activity of the SaeRS two-component system (TCS). Deletion of the cardiolipin synthase genes, *cls1*, and *cls2*, reduces the basal activity of SaeRS and other TCSs. Cardiolipin is an indispensable requisite for Sae activation mediated by human neutrophil peptides (HNPs) in the stationary growth phase but not mandatory for Sae induction in the exponential growth phase. Ectopic expression with *cls2*, but not with *cls1*, in the *cls1 cls2* double mutant fully restores Sae activity. Elimination of cardiolipin from the membranes results in decreased kinase activity of the sensor protein SaeS. Purified SaeS protein directly binds to cardiolipin as well as phosphatidylglycerol. A strain lacking *cls2* or *cls1cls2* renders *S. aureus* less cytotoxic to human neutrophils and less virulent in a mouse model of infection. Our findings suggest that cardiolipin enables a pathogen to confer virulence by modulating the kinase activity of SaeS and other sensor kinases upon infection.

## Introduction

*Staphylococcus aureus* is a Gram-positive pathogenic bacterium and a major driver of antibiotic resistance and infections (1, 2). The bacterium causes a variety of diseases ranging from soft-tissue infections to life-threatening diseases, such as endocarditis, sepsis, toxic shock syndrome, and necrotizing pneumonia (3). While the number of both hospital-acquired and community-associated *S. aureus* infections has increased over the past decades, the treatment of the bacterial infections has become more difficult due to the emergence of multi-drug resistant strains (4–6). *S. aureus* utilizes multiple regulatory pathways including two-component signal transduction systems (TCSs) to control the expression of virulence factors, which in turn contributes to the pathogenesis of the bacterium (7, 8). The TCSs are a tunable genetic device that employs reversible protein phosphorylation to modify a gene regulatory circuit in response to changing environmental conditions. Of the 16 TCSs in *S. aureus*, the SaeRS TCS governs the expression of over 20 critical virulence factors (e.g., α-toxin, nuclease, coagulase, proteases, leukocidins) that confer staphylococcal virulence (7). The system is composed of the sensor histidine kinase SaeS and its cognate response regulator SaeR (7, 9). Upon exposure to host-derived signals, such as human neutrophil peptides (HNPs) (7, 9), SaeS autophosphorylates the conserved histidine residue and subsequently transfers the phosphoryl group to the conserved aspartate residue of SaeR. Then, phosphorylated SaeR (SaeR-P) binds to the upstream region of its target genes and promotes their transcription (7).

Fine-tuning of the membrane composition is essential for bacterial survival and adaptation in response to the growth phase or various environmental stresses such as temperature, organic solvents, nutrient starvation, or pH (10–17). The membrane phospholipids in *S. aureus* consists of the three polar phospholipids: phosphatidylglycerol (PG), lysyl-phosphatidylglycerol (L-PG), and cardiolipin (CL) (18). In actively growing cells, PG is the predominant phospholipid, whereas CL is the major phospholipid in stationary growth phase, in which two PG molecules are converted into CL and glycerol by cardiolipin synthase (12, 19). *S. aureus* has two CL synthase genes, *cls1* and *cls2* (12, 20). *cls2* encodes the major CL synthase accountable for CL accumulation in stationary phase and following phagocytosis by neutrophils (12, 20, 21). Hence, Cls1 is responsible for CL synthesis under acid stress condition (19, 20).

In *S. aureus*, except for NreB, all other 15 TCS sensor kinases are membrane proteins with at least two transmembrane domains. For this reason, we hypothesized that their activity would be affected by changes in the membrane composition. Indeed, recently, we and others showed that changes in membrane phospholipids (*e*.*g*., branched-chain fatty acids or a pool of intracellular and exogenous fatty acids) impact the activity of the SaeRS as well as a set of other TCSs (22–24). As the CL content changes during *S. aureus* growth, here we examined whether the CL content changes impacts the activation of Sae.

## Results

### Cardiolipin is required for the full activity of Sae

To investigate the role of cardiolipin (CL) in the SaeRS TCS activity, we deleted the CL synthase genes, *cls1* and *cls2*, in *S. aureus* USA300. Deletion of *cls1* did not affect the level of CL noticeably, whereas deletion of *cls2* did, and almost no CL was detected in the *cls1cls2* double mutant (Fig. 1A), confirming the previous report that Cls2 is the major CL synthase in *S. aureus* (12). The P1 promoter of the *sae* operon has two SaeR binding sites and is an excellent indicator for the Sae activity (7, 25). To measure the Sae activity in those deletion mutants, we introduced the P1-*gfp* reporter plasmid, pYJ-*sae*P1-*gfp*, into the *cls1, cls2*, and *cls1cls2* deletion mutants and measured GFP expression. The P1 activity was marginally reduced in the *cls1* mutant, whereas it was greatly reduced in the *cls2* and *cls1cls2* mutants (Fig. 1B), exhibiting the similar trends as with the CL levels. Tsai *et al*. reported that CL levels increase toward the stationary growth phase (20). Since the P1 activity seems to be activated by CL, we speculated that the P1 activity would also be higher at the stationary growth phase. Indeed, when the P1 activity was measured throughout the bacterial growth phases, the P1 activity in the WT strain also increased toward the stationary growth phase and kept increasing (Fig. 1C. solid lines). On the other hand, all *cls* mutants exhibited reduced P1 activity during cell growth throughout the growth phase (Fig. 1C). As expected, almost no Sae activity was detected in *sae*-deletion strains (Fig. 1C). Also, no growth defect was observed in all *cls* mutants (Fig. 1C. dotted lines).

**Fig. 1.**
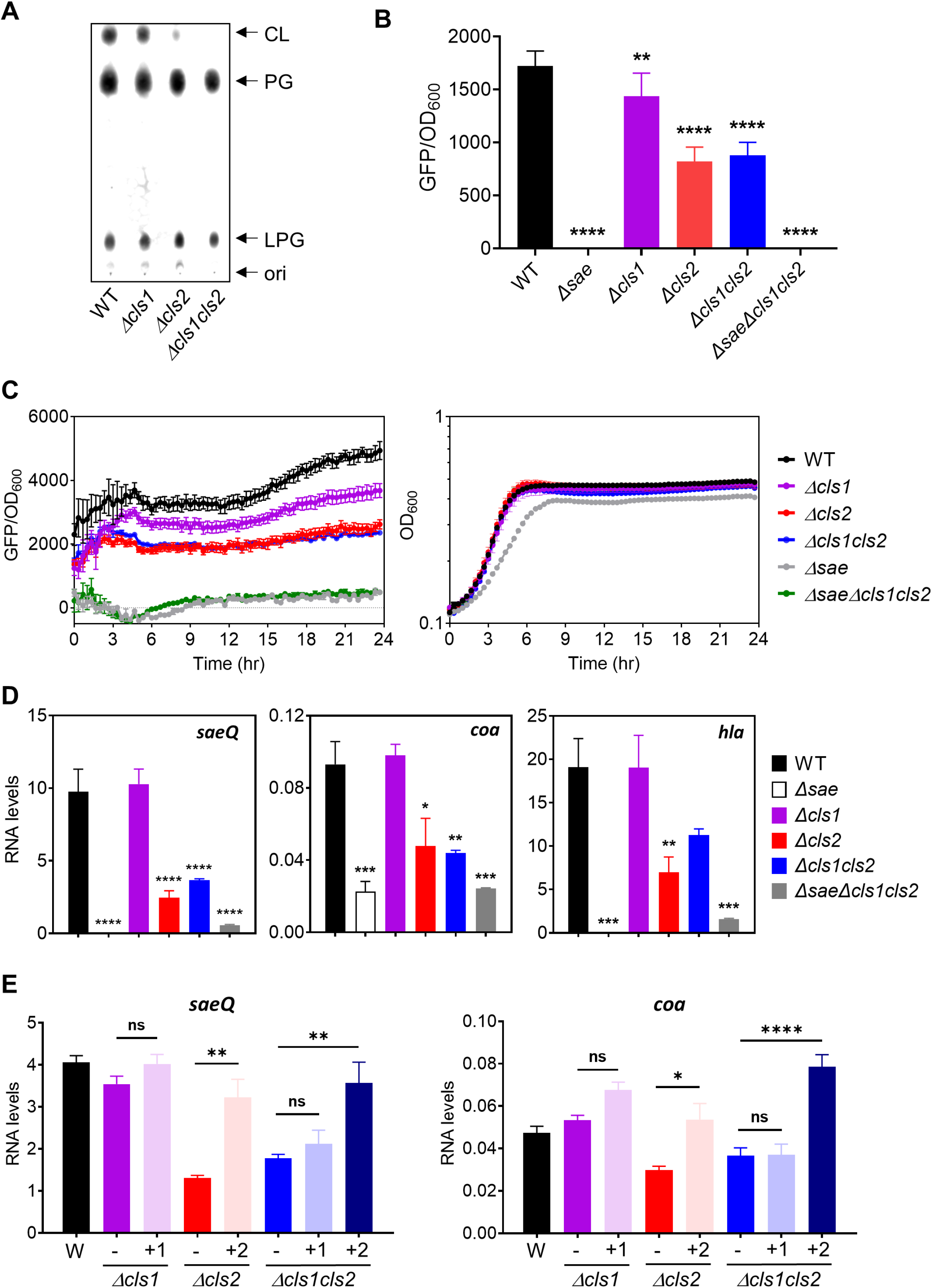
Inactivation of the *cls* genes reduces the basal expression of the Sae TCS system. (A) Phospholipid composition of USA300 and its isogenic *cls* mutants. Bacteria were harvested at 24 hr (stationary phase) of growth in TSB at 37°C, and lipids were extracted and resolved by thin layer chromatography (TLS). CL: Cardiolipin. PG: Phosphatidylglycerol. LPG: lysyl-phosphatidylglycerol. ori: origin. The TLCs are representative of three independent experiments. (B) Cells were grown overnight in TSB and *sae*P1-GFP activity in the indicated strains harboring a *sae*P1-*gfp* reporter fusion (pYJ-*sae*P1-*gfp*) was measured. All experiments were repeated at least three times with similar results observed. Error bars represents the standard error of the mean. Unless indicated otherwise, GFP expression was compared with that of the strain carrying wild type SaeS, and statistical significance was assessed by a one-way ANOVA with Dunnett’s *post* test, relative to wild type (WT). **, p<0.0014; ****, p<0.0001. (C) Wild-type (WT), *Δsae, Δcls1, Δcls2, Δcls1cls2* and *ΔsaeΔcls1cls2* strains harboring the *sae*P1-*gfp* reporter fusion (pYJ-*sae*P1-gfp) were grown in TSB at 37°C and *sae*P1 expression was monitored for 24 h. Relative fluorescence units (GFUs) (left) and cell density (optical density at 600 nm [OD_600_]) (right) and were measured. Dotted lines, cell OD_600_; solid lines, GFP/OD_600_. All data correspond to the mean ± SEM from at least three independent experiments. (D) mRNA levels of the *saeQ, coa* and *hla* genes produced by the indicated strains grown in TSB for 6 hr. mRNA levels were normalized to those of the *gyrB* gene. Statistical significance was assessed by a one-way ANOVA with Dunnett’s *post* test, relative to WT. *, P<0.0139; **, p<0.0081; ***, p<0.0006; ****, p<0.0001. (E) Ectopic expression with the *cls2* gene restores Sae activity. mRNA levels of genes specific to other two-component systems. RNA was extracted from strains grown in TSB for 6 hr and used for RT-qPCR. mRNA levels were normalized to those of the *gyrB* gene. Statistical significance was assessed by a one-way ANOVA with Dunnett’s *post* test, relative to wild type (WT). −, the corresponding *cls* mutant; +1, pOS1-*cls1*; +2, pOS1-*cls2*. *, P<0.0177; **, p<0.0057; ****, p<0.0001; ns, not significant.

To further confirm the positive role of CL in the Sae activity, we measured the effect of CL on the transcript levels of three well-known Sae regulated genes: *saeQ, coa*, and *hla* (7). As can be seen, the deletion of *cls2* significantly decreased the transcript levels of all three genes (Fig. 1D). Ectopic expression with *cls2* under the control of the native *cls* promoter using a multicopy plasmid pOS1 (26) fully restored Sae activity of the *sae* target genes in the *cls2* and *cls1 cls2* mutant strains, but not with *cls1* (Fig. 1E). These results demonstrate that CL is required for the full activity of Sae.

### Cardiolipin is not required for the Sae activation by human neutrophil peptides

Human neutrophil peptides (e.g., HNP1) activate the SaeRS TC system (9, 27). We next sought to examine whether cardiolipin is required forSae activation by HNP1. USA300 WT and the *cls* mutants carrying pYJ-*sae*P1-*gfp* were grown in TSB with or without HNP1 (5 μg/ml), and P1 promoter activity was measured at the exponential (4 h growth) and the stationary growth phase (24 h growth). At the exponential growth phase, basal Sae activity was reduced, particularly, in the *cls2* mutant; however, Sae activity was activated by HNP1 in all the mutants to levels similar to that of WT (Fig. 2A). Similar results were observed in the cells grown to the stationary growth phase, although the Sae activation levels were somewhat lower in the cells lacking both *cls1cls2* than the WT level (Fig. 2B), indicating that CL plays a minor role during HNP1-mediated Sae activation.

**Fig. 2.**
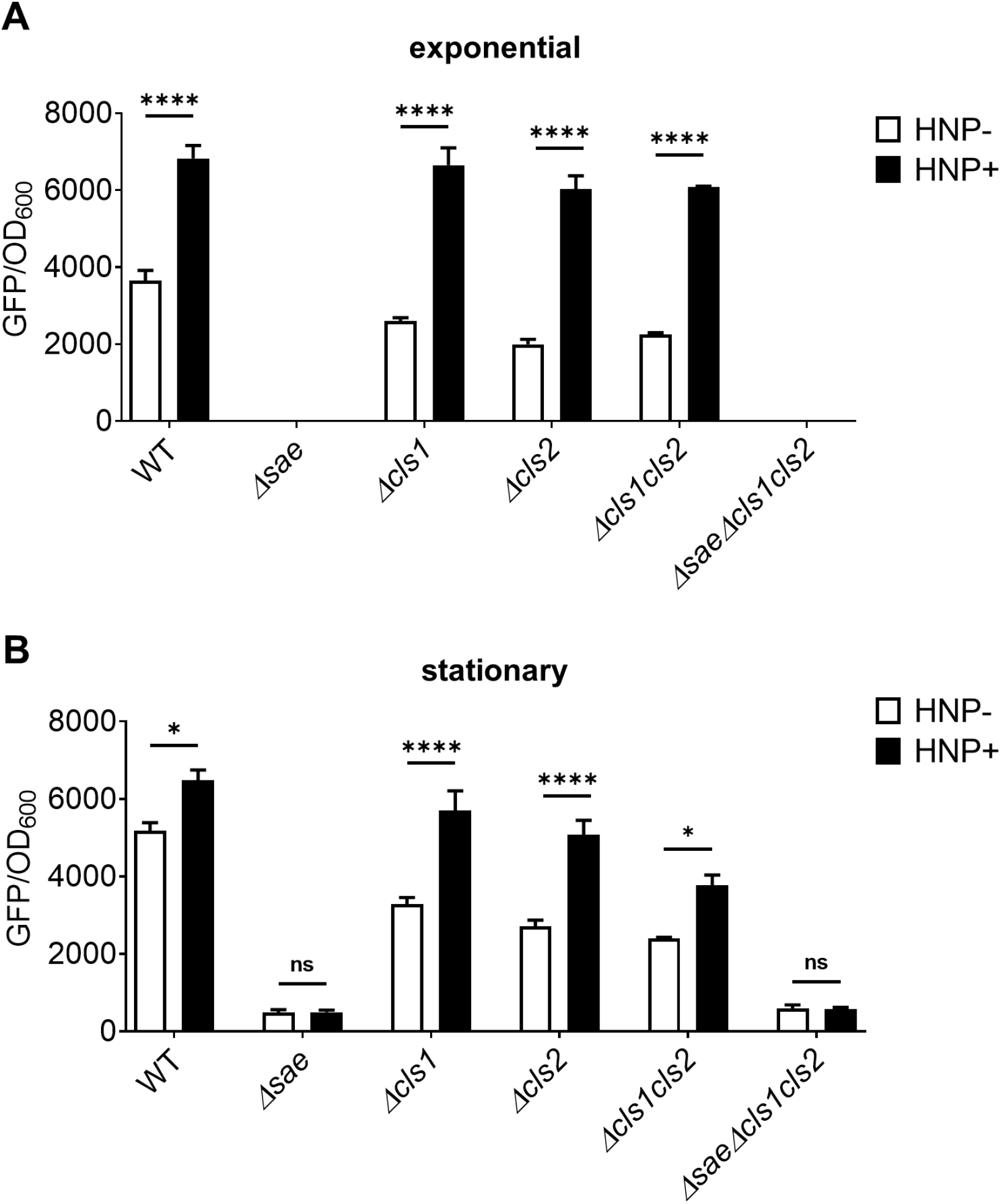
Cardiolipin is dispensable to activate the Sae system upon exposure to HNPs. Cells were grown in fresh TSB for 2hr and further incubated in the absence or presence of HNP1. *sae*P1 promoter activity was measured in the exponential phase (A) and stationary phase (B). The levels of GFP expression were normalized to OD_600_ value. All data correspond to the mean ± SEM from three independent experiments. A two-way ANOVA with Tukey’s multiple comparisons was used for statistical analysis between HNP1 (-) and HNP1 (+). *, p<0.0238, ****, p<0.0001.

### Cardiolipin-deficient membranes have reduced SaeS kinase activity

Next, we investigated how CL positively affects Sae activity. As with other two-component systems, Sae activity is determined by SaeS kinase activity (i.e., the autokinase and phosphotransferase activities) (7, 27). Since the P1 promoter activity was reduced in the *cls* mutant (Figs 1 and 2), we suspected that CL is required for SaeS kinase activity. To test this notion, we isolated membrane vesicles from USA300 WT and three mutants, *cls1, cls2*, and *cls1 cls2*, grown to the early stationary growth phase. The membrane vesicles were then used as the source of SaeS in a kinase assay *in vitro*. The SaeS kinase activity was determined by measuring the levels of phosphorylated recombinant SaeR protein that was added in the reaction with [γ-^32^P] ATP. SaeR-P was normalized by the level of SaeS protein in the membranes determined by Western blot analysis and densitometry (Fig. S1). The levels of SaeR-P catalyzed by SaeS in the *cls2 or cls1cls2* mutant membranes were about two-fold lower than that from wild-type membranes (Fig. 3A and 3B), agreeing with the *sae*P1 promoter activity (Fig. 1B). Based on these results, we concluded that CL is required for SaeS kinase activity.

**Fig. 3.**
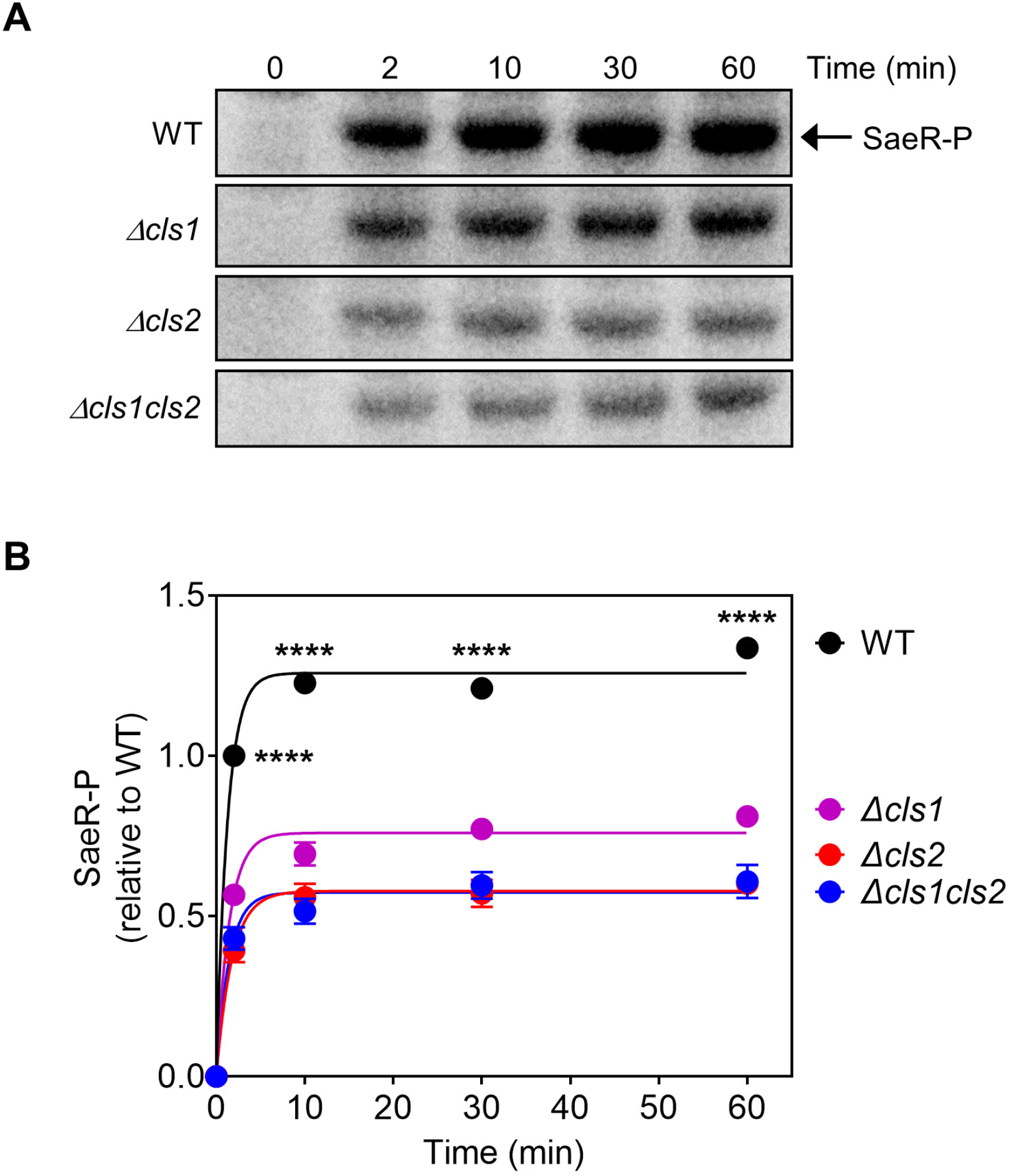
Lack of cardiolipin results in decreased SaeS kinase activity. **(A)** Levels of SaeR-P after incubation of recombinant His_6_-SaeR with the wild-type (WT), *cls1, cls2*, or *cls1cl2* mutant membrane vesicles from USA300 in the presence of [γ-^32^P] ATP as a source of ATP (Top). Levels of SaeR-P were normalized by the level of the SaeS protein in the membrane, which was determined by Western blot analysis (Fig. S1). (B) The graph depicts the levels of SaeR-P at the indicated time point relative to the SaeR-P at the 2 min time point (not shown) in the wild-type membranes using densitometry analysis (bottom). All data correspond to the mean ± SEM from three independent experiments. For statistical analyses, a two-way ANOVA with Tukey’s multiple comparisons was used for statistical analysis comparing each SaeR-P level between wild-type (WT) and *cls* mutant at the indicated time points. ****, p<0.0001.

### SaeS binds to cardiolipin

Some bacterial membrane proteins interact with membrane phospholipids, including CL, which are required for various functions such as stability, oligomeric state, or protein folding (28–31). Since SaeS has two transmembrane domains and requires CL for its kinase activity, it is possible that SaeS can directly interact with CL. To test this possibility, we purified a maltose-binding protein (MBP)-SaeS, which has a His_6_-epitope tag at the N-terminus and retains its kinase activity *in vitro* (27). The recombinant MBP-His_6_-SaeS protein was incubated with a membrane strip prespotted with 15 different lipid species (Echelon Biosciences). The bound MBP-His_6_-SaeS protein was detected using an anti-SaeS or anti-His_6_ antibody (Fig. 4). For a control, a His-tagged MBP (MBP-His_6_) protein was used. As shown, MBP-His_6_-SaeS bound not only CL but also PG, the main phospholipid components in *S. aureus* (Fig. 3). Unexpectedly, it showed strong affinity to sulfatide, a sulfolipid, which is only found in eukaryotic cell membranes. In addition, it exhibited modest binding to phosphatidylserine (PE) (Fig. 4). Since neither sulfatide nor phosphatidylserine is present in *S. aureus* cell membrane, their binding to SaeS is not physiologically relevant. The control protein MBP-His_6_ did not bind to any of the lipids tested (Fig. 4). These results suggest that CL enhances SaeS kinase activity via direct binding.

**Fig. 4.**
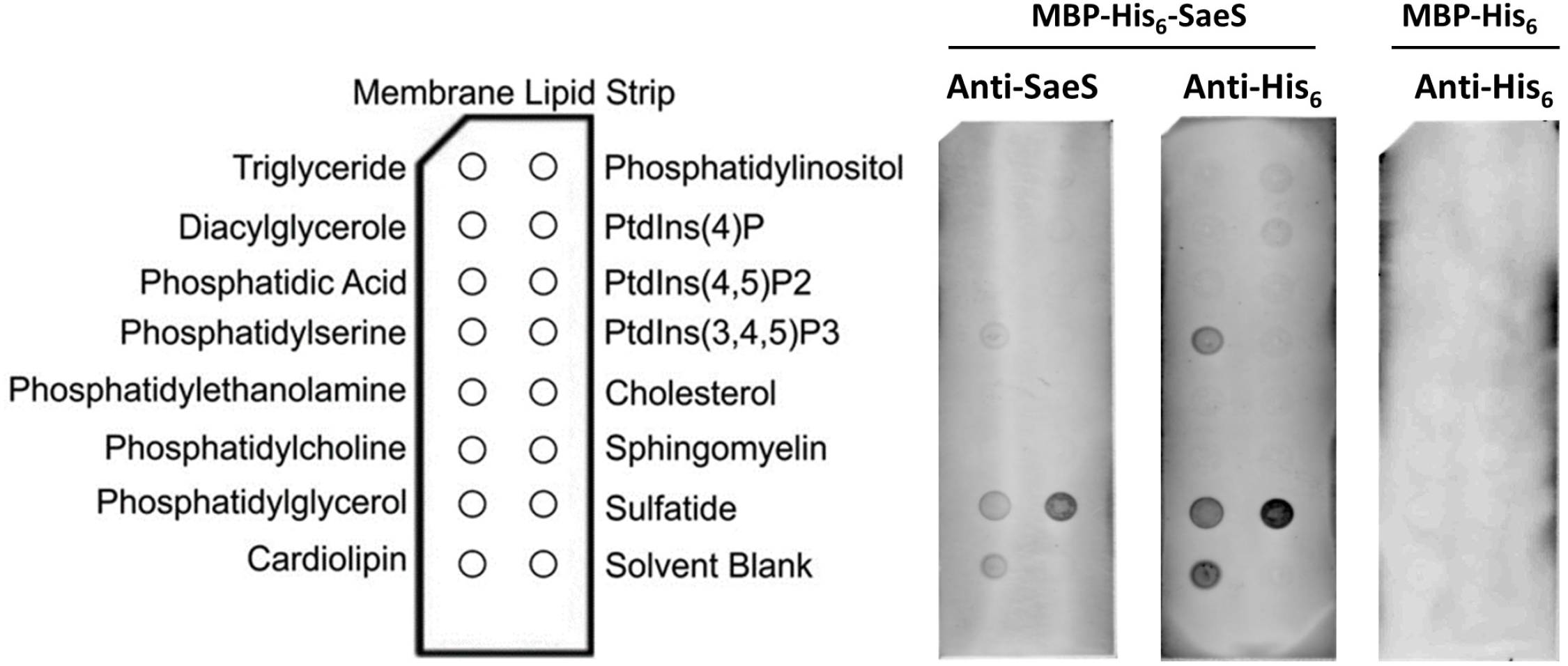
SaeS binds to cardiolipin. Membrane lipid strips prespotted with 15 different lipid species were incubated with 2.5 μM of the purified MBS-His_6_-SaeS and MBP-His_6_ proteins. The bound SaeS proteins were detected using anti-SaeS and anti-His_6_ antibody and StarBright Blue 700 Goat Anti-Mouse. The image is representative of at least three independent experiments.

### The transcription of seven TCSs is also affected by CL

*S. aureus* harbors 16 TCSs. With the exception of NreB (oxygen-responsive, cytoplasmic sensor kinase), all sensor kinases are situated in the membrane (8). To investigate whether CL alters the activities of other TCSs, we took advantage of the fact that TCSs exhbit feedback autoregulation of the sensor kinase and response regulator genes (32) and measured the transcript abundance of sensor kinases of all 16 TCSs in the WT and the *cls* mutant strains. As shown, transcription of three additional sensor kinases, *arlS, hptS*, and *lytS* showed a similar pattern to that of *saeS*; it was reduced only when *cls2* was deleted (Fig. 5). Intriguingly, however, transcription of *braS, desk, graS*, and *vraS* was reduced in all *cls* mutants (Fig. 5). The transcript abundances of the remaining sensor kinase genes including *nreB* (i.e., *agrC, airS, hssS, kdpD, nreB, phoR, ssrB*, and *walR*) were not significantly affected by the *cls* mutations (Fig. S2). These results suggest that CL plays an important role in maintaining the proper activity of eight TCSs in *S. aureus*.

**Fig. 5.**
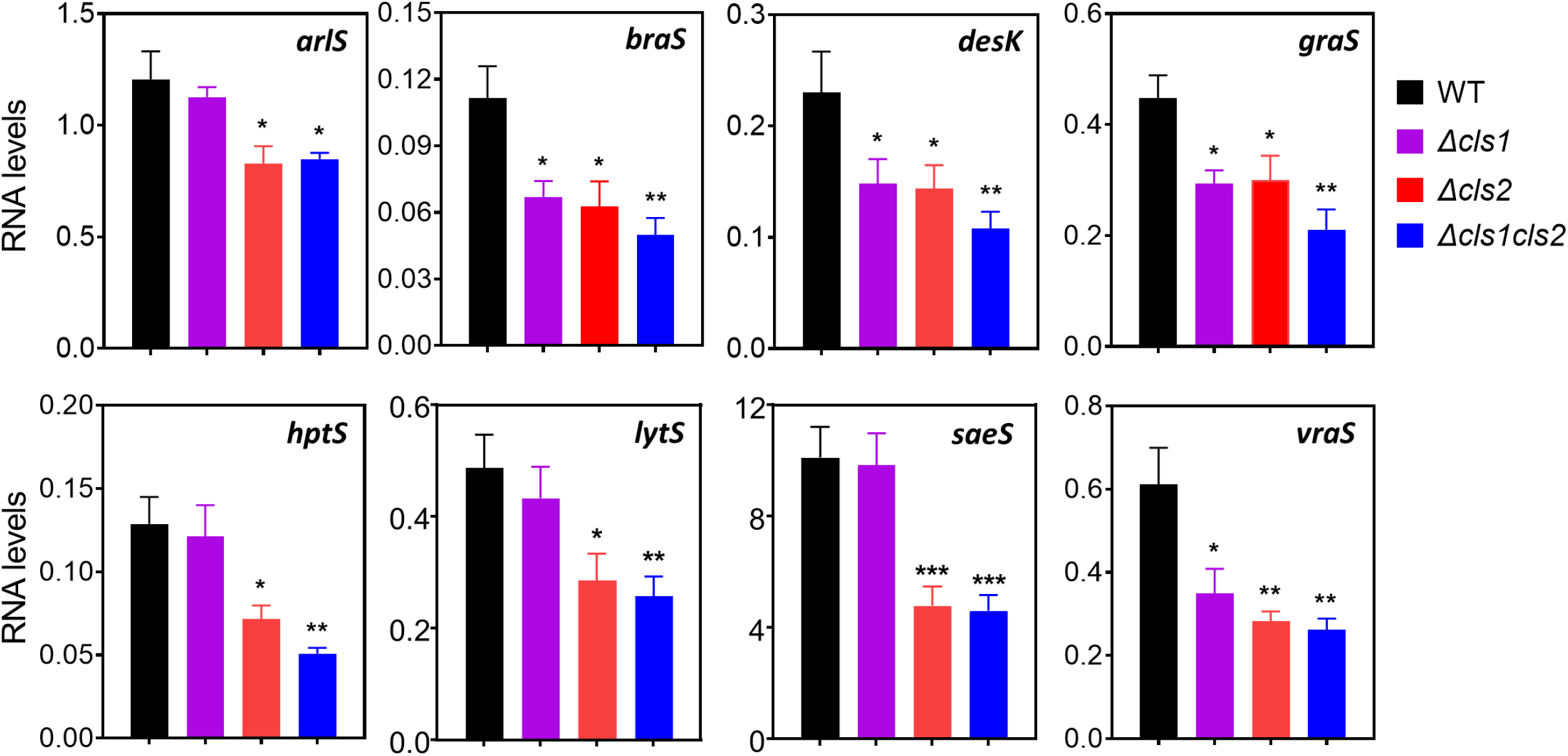
Cardiolipin is required for the basal expression of a subset of TCSs. mRNA levels of genes specific to eight of staphylococcal TCSs. RNA was extracted from the wild-type (WT), *Δcls1, Δcls2*, and *Δcls1cls2* strains grown in TSB for 6 h and used for reverse transcription quantitative PCR (RT-qPCR). mRNA levels were normalized to those of the *gyrB* gene. A one-way ANOVA with Dunnett’s *post* test was used for statistical analysis comparing transcript abundance between WT and the mutants. *, p<0.0454; **, <0.0091; ***, p<0.0008. All data correspond to the mean ± SEM from at least three independent experiments, which produced similar results.

### *cls2* is critical for the virulence of *S. aureus*

Lack of *sae* or decreased SaeS kinase activity greatly reduces staphylococcal virulence in various host environments (27, 33). Also, seven more TCSs appear to require CL for their full activity (Fig. 5). To investigate the contribution of CL to staphylococcal virulence, we examined the cytotoxicity of culture supernatants of the WT and the three *cls* mutant strains toward human PMNs (hPMNs). The test strains were grown in TSB to the post-exponential growth phase (5 h and 8 h), and culture supernatants were obtained and mixed with purified hPMNs. As expected, the culture supernatant of the *sae*-deletion mutant showed minimal cytotoxicity (Fig. 6A). On the other hand, the culture supernatants of the three *cls* mutants showed modestly reduced cytotoxicity toward human hPMNs compared to WT USA300 (Fig. 6A), reflecting the reduced Sae activity shown in Fig. 1 and 2. The combination of deletion of *sae* and *cls1 cls2* exhibited similar cytotoxicity to that observed in the *sae* mutant (Fig. 6A).

**Fig. 6.**
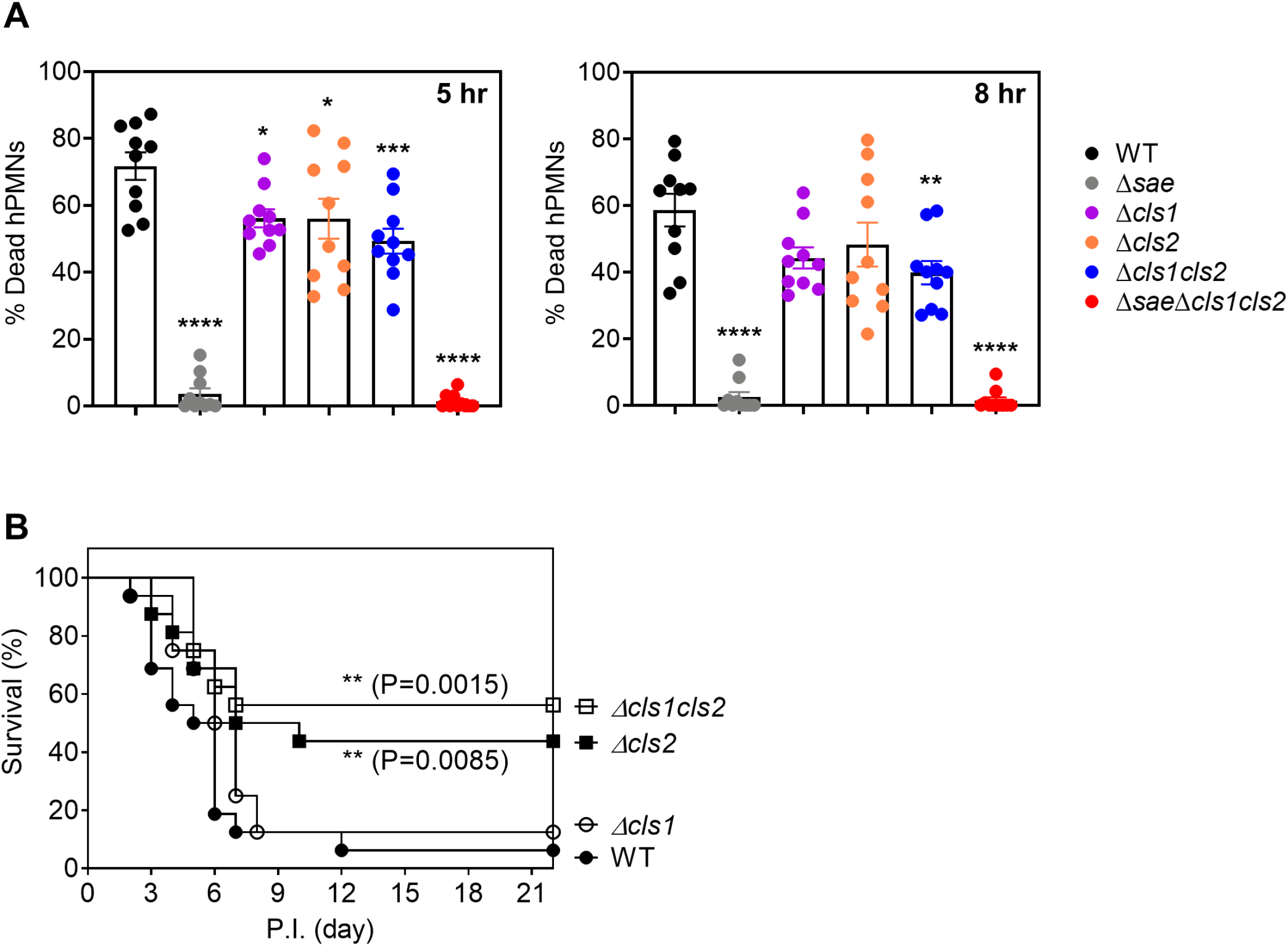
Lack of cardiolipin reduces staphylococcal virulence with respect to human neutrophils and mice. (**A)** Primary human PMNs (hPMNs) were intoxicated with 2.5% supernatants obtained from the wild-type (WT), *Δsae, Δcls1, Δcls2, Δcls1cls2*, and *ΔsaeΔclscls2* mutants at either 5 h or 8 h following subculture. hPMN viability was measured by CellTiter 96 (Promega). Data show mean ± SEM of PMNs isolated from 10 independent blood donors. Statistical significance was assessed by a one-way ANOVA with Dunnett’s *post* test, relative to wild-type (WT). *, p<0.0143; ***, p<0.0002; ****, p<0.0001. (B) C57BL/6 mice (N=16; male=8 and female=8) were retro-orbitally infected with 2.5 × 10^7^ CFU of wild-type (WT), *Δcls1, Δcls2*, and *Δcls1cls2* mutant *S. aureus*. Survival of mice was monitored daily. The differences in the survival were assessed using Log-rank (Mantel-Cox) test.

To further investigate the contribution of CL to staphylococcal virulence, we infected mice with the WT or the three *cls* mutant strains via retro-orbital injection and observed the infected mice for 22 days. The mice infected with WT or the *cls1* mutant exhibited 95% and 90% mortality, respectively. On the other hand, mice infected with *cls2* mutant or the *cls1* and *cls2* double mutant exhibited only 50% and 40% mortality, respectively (Fig 6B). Considering the correlation between the CL levels in the cell membranes (Fig. 1A) and the killing of mice (Fig. 6B), we concluded that CL is required for the virulence of *S. aureus*.

## Discussion

Membrane proteins, including most bacterial TCS sensor kinases, are embedded in the cytoplasmic membrane and expected to interact with membrane lipids. In this study, we showed that CL, a major phospholipid of *S. aureus* cytoplasmic membrane, directly interacts with the SaeS sensor kinase of the Sae TCS (Fig. 4) and positively affects its kinase activity (Fig. 3A). CL was also required for bacterial virulence (Fig. 6). However, the positive effect of CL was not limited to SaeS; the transcript abundance of seven of other TCS sensor kinases was also reduced in the *cls1* and *cls2* mutants. Therefore, the alteration in the CL content or changes in membrane composition in general might impact the activities of other membrane proteins, if not all. Cosidering the fact that bacterial membrane composition could be altered during growth, it is plausable that membrane changes might, at least in part, contribute to some of the growth-dependent alterations in bacterial physiology and gene regulation.

Structure function analysis indicates that the head group of phospholipids is critical for binding to SaeS (Fig. 4). Phospholipids with serine or glycerol in the head group bind to SaeS, whereas no significant SaeS binding was observed for phospholipids with ethanolamine or choline. The critical role of the head group in SaeS binding also suggests that SaeS interacts with phopsholipids through the domains situated close to the surface of the membrane. Since, in *S. aureus*, all three major phospholipids (*i*.*e*., PG, CL, and Lysyl-PG) contain glycerol in their head group, it is likely that SaeS directly interacts with all of those phospholipids. However, the positive impact of CL on the SaeS kinase activity implies that CL induces a conformational change that is distinct from that of PG.

A prototypical sensor histidine kinase contains a long extracellular sensory domain that directly senses external stimuli (8, 34). However, a subset of sensor kinases called intramembrane-sensing HKs (IM-HKs) lacks a sensory domain, and their transmembrane domains are connected by a short linker peptide (<25 amino acids) (35). In *S. aureus*, there are four TCSs carrying IM-HKs: SaeRS, GraRS, BraRS, and VraSR. Intriguingly, transcription of those IM-HKs were all reduced in *cls2* or *cls1*and *cls2* mutants (Fig. 5). Although the linker peptides are not expected to directly bind to their stimuli, substitution mutations in the peptide could alter the kinase activity of IM-HKs (27), suggesting that the conformational changes in the linker peptides can be transmitted to the cytoplasmic domain and alter enzymatic activities of IM-HKs. Since the linker peptides are likely in a close contact with the surface of the cytoplasmic membrane (i.e., the head groups of phospholipids), it is possible that CL’s positive impact on the SaeS kinase activity is through interaction with the linker peptide. Recently, we showed that branched-chain fatty acid in membrane phospholipids increases the Sae activity (23). Since fatty acids are not exposed to outside of the membrane, CL and fatty acids might affect the SaeS activity in a different mechanism. However, it remains to be determined whether the other IM-HKs (i.e., GraS, BraS, and VraS) also bind to CL, and if they do, the binding is via the linker peptide or other domains close to the surface of the membrane.

Unexpectedly, we found that SaeS strongly interacts with sulfatide (Fig. 4). Sulfatide is 3-*O*-sulfogalactosylceramide, the first sulfoglycolipid isolated from the human brain (36). Although sulfatide does not exist in *S. aureus* membrane, sulfatide treatment is known to improve the survival rate of mice with *S. aureus* lethal sepsis by downregulating systemic inflammation without affecting bacterial loads in blood, liver, and kidneys of the infected mice (37, 38). Since the SaeRS TCS activates the production of over 20 virulence factors, it might be possible that sulfatide treatment dampens the Sae activity and diminishes a cytokine storm induced by the bacteria, resulting in a decrease of lethality in the early stage of sepsis. This conjecture also implies that the interaction with sulfatide would decrease the SaeS kinase activity.

## MATERIALS AND METHODS

### Ethics statement

Leukopaks were obtained from de-identified healthy adult donors with informed consent from the New York Blood Center and were used for the isolation of neutrophils. De-identified samples are exempted from the ethics approval requirements by the NYULH Institutional Review Board. All animal experiments were performed in accordance with the Guide for the Care and Use of Laboratory Animals of NIH. The animal protocol was approved by the IUSM NW IACUC (#NW-48). Every effort was made to minimize the suffering of the animals.

### Bacterial Strains, Plasmids and Culture Conditions

The bacterial strains and plasmids used in this study are listed in Table S1. *Escherichia coli* was grown in LB (39), while *S. aureus* was cultured in tryptic soy broth (TSB) containing 0.25% (wt/vol) dextrose (BD Biosciences). For transduction of marked alleles and plasmids, heart infusion broth (HIB) supplemented with 5 mM CaCl_2_ was used. When necessary, antibiotics were added to the growth media at the following concentrations: ampicillin (Amp), 100 μg/ml; erythromycin (Erm), 10 μg/ml; and chloramphenicol (Chl), 5 μg/ml.

### DNA Manipulation

Unless stated otherwise, all restriction enzymes and DNA modification enzymes were purchased from New England Biolabs. For PCR amplification, the Phusion DNA polymerase (New England Biolabs) was used. Plasmids and genomic DNA were extracted with Zippy™ plasmid miniprep kit (Zymo Research) and GenElute™ Bacterial Genomic DNA kit (Sigma), respectively, according to the manufacturer’s instruction. Plasmid DNA was introduced into *E. coli* by the method of Hanahan and Meselson (40) and into *S. aureus* RN4220 by electroporation with a gene pulser (Bio-Rad). All plasmids were verified using restriction digestion or Sanger sequencing. Subsequent transduction of the plasmids into target strains of *S. aureus* was carried out with φ85 (41).

### Construction of clean deletions *Δcls1, Δcls1*, and *Δcls1 cls2* strains

To construct *Δcls1* and *Δcls2* strains, DNA sequences 1 kb upstream and downstream of *cls1* or *cls2* were PCR-amplified using the primer pairs P2551/2552 and P2553/2554 (Table S2). The target vector pIMAY (42) was also PCR-amplified with primer pairs P1986/1987 (Table S2). The PCR products were assembled by the ligation independent cloning method (43). First, the insert DNA and vector PCR products were treated with T4 DNA polymerase for 30 min at room temperature. Then the DNA fragments were mixed and incubated at 37°C for 30 min and transformed into *E. coli*. The pIMAY containing the *cls1* or *cls2* deletion cassette was isolated and electroporated into RN4220. Subsequently, the plasmid was moved into USA300 strain by ϕ85-mediated transduction. Deletion of *cls1* and *cls2* was carried out by following the procedures previously reported (44) and verified by PCR amplification of the gene locus followed by DNA sequencing.

### Construction of plasmids

For complementation in the *cls* mutant, pOS1-*cls1* and pOS1-*cls2* were constructed in pOS1(26) using primers P2434/P2435 and P2436/P2437, respectively. The fragments were digested with *Bam*HI/*Sal*I and ligated to the same site of pOS1 and the ligated plasmids were inserted into *E. coli* DH5α. All plasmids were confirmed by PCR in addition to Sanger sequencing. The recombinant plasmids were electroporated into *S. aureus* RN4220, from which the plasmids were further transduced into the *Δcls1, Δcls2* or *Δcls1cls2* strains with Φ85.

pET22b-*saeR* was constructed in pET22b (Novagen) using primers P2954/P2955. The fragments were digested with *Nde*I/*Hindl*III and ligated to the same site of pET22b and the ligated plasmids were inserted into *E. coli* DH5α. The resulting plasmid was confirmed by PCR in addition to Sanger sequencing and transformed into BL21 (DE3).

### GFP Reporter Assays

We performed the GFP reporter assays using a microplate reader (Enspire, PerkinElmer). For the *sae*P1-GFP assays, overnight cultures of the test strains were diluted 1:100 into fresh TSB (2 ml) and incubated at 37°C with shaking. After 2 hr incubation, each culture was divided into two (1 ml each), and human neutrophil peptide 1 (HNP1, 5 μg/ml) was added to one sample and further incubated at 37°C for 2 hr. At 4 hr and 24 h post incubation, 100 μl of cell cultures from at least three biological replates were placed in a black 96-well plate in duplicate, and the fluorescence (485 nm excitation, 538 nm emission) was measured in a microplate reader (Enspire, Perkin Elmer). To measure the *sae*P1 activity during cell culture for 24 hr, cultures (2 ml) inoculated from single colonies (at least three biological replates) were grown 16 hours, normalized by cell OD_600_ (final OD=0.4) and diluted 1: 100 into 150 μl of fresh TSB media in flat-bottomed 96-well plates (PerkinElmer). The cultures were overlaid with 100 μl of mineral oil (Sigma) to prevent evaporation. The cultures were grown in an Enspire multiwell fluorimeter (PerkinElmer) set at 37°C and assayed with an automatically repeating protocol of shaking (1 mm orbital, fast speed, 30 s), fluorescence readings, and absorbance measurements at 15 min intervals for 24 hr. Background fluorescence of cells bearing a promotorless GFP vector was subtracted. The fluorescence was normalized by OD_600_.

### Membrane preparation

Membrane vesicles were prepared as described previously with a slight modification (27). The wild type and *cls* mutants were grown for 6 hr in TSB. Cell pellets were frozen and resuspended in membrane lysis buffer (100 mM Tris pH 8.0, 100 mM NaCl, 10 mM MgCl_2_) containing 40 μg/mL lysostaphin and incubated for 20 minutes at 37°C. Cells were disrupted by sonication (FisherBrand Model 120 Sonic Dismembrator). After removing cell debris, membranes were recovered via ultracentrifugation at 82,700 × g for 30 minutes (Optima XPN-90 Ultracentrifuge, Beckman). Finally, the pellet was resuspended in TKMG buffer (50 mM Tris pH 8.0, 50 mM KCl, 1 mM MgCl_2_, 25% glycerol) and used as the inverted membrane vesicles in this study. The protein concentration was determined using a BCA protein assay kit (Pierce) with bovine serum albumin as a standard.

### Western Blotting

100 μg of membranes were subjected to SDS-PAGE and proteins were transferred to polyvinylidene difluoride (PVDF) membranes (Cytiva). Membranes were blocked with 10% [w/v] skim milk and incubated with SaeS antibody (1:1000) for 1 h at room temperature. Membranes were washed with TBST and incubated with StarBright Blue 700 Goat anti-rabbit IgG (1:3500; Bio-Rad) for 1 hour. After a brief wash in TBST, the signals were visualized using an Amersham ImageQuant 800. The densities (mean intensity per unit area) of the SaeS and SaeR protein bands were determined by quantification with Multi Gauge software (Fuji Film).

### Kinase Activity Assays

Three to five hundred microgram of membrane vesicles were mixed with 10 μM of the SaeR-His_6_ protein in 1×TKM buffer (50 mM Tris pH 8.0, 50 mM KCl, 1 mM MgCl_2_) at room temperature for 5 min. The reaction was started by incubating with 0.1 mM ATP containing 20 μCi of [γ-^32^P] ATP (3000 Ci/mmol; Perkin Elmer) containing 1 mM of cold ATP at room temperature. A 9 μl aliquot was mixed with 5 × SDS sample buffer at different time points to stop the reaction. The phosphorylated SaeR-His_6_ proteins were separated by 12% Bis-Tris SDS-PAGE and determined by quantifying the [^32^P]-labeled species using an Amersham Typhoon Biomolecular Imager and a phosphor imaging plate (Fuji Film) followed by quantification with Multi Gauge software V 3.0 (Fuji Film). The data were fitted using nonlinear regression to a one phase exponential association (Prism 9, GraphPad). All data correspond to the mean ± SEM from three independent experiments.

### MBP-SaeS and SaeR purification

The MBP-His_6_-SaeS and SaeR-His_6_ proteins were overproduced in *E. coli* BL21 (DE3) harboring plasmids pMCSG19-saeS (27) or pET22b-saeR, respectively. The MBP-His_6_-SaeS protein was purified as described before (27). To purify SaeR-His_6_, overnight cultures were inoculated into fresh LB broth, and the proteins were expressed by the addition of 1 mM of isopropyl-1-thio-β-D-galactopyranoside (IPTG) to the culture, and further incubated at 37°C for 6 h. The proteins were purified with Ni-column chromatography (Qiagen) by following the manufacturer’s recommendations.

The purified MBP-His_6_-SaeS and SaeR-His_6_ were exchanged and concentrated with Amicon (Ultracell-30 for MBP-His_6_-SaeS, Ultracell-15 for SaeR-His_6_; Millipore) in TKM buffer for SaeS (50 mM Tris-Cl [pH 8.0], 50 mM KCl, 1 mM MgCl_2_) or TBS buffer for SaeR (10 mM Tris-HCl [pH 7.5], 138 mM NaCl, 2.7 mM KCl), respectively, and then in TKM or TBS buffer containing 25% [v/v] glycerol. Protein concentration was determined by the bicinchoninic acid assay (Bio-Rad), and the purified proteins were stored at −80°C until used.

### Lipid extraction and thin-layer chromatography

Cells equivalent to 2 × 10^8^ CFU were collected at stationary phase (24h culture in TSB) and normalized to 1 ml of 2.0 OD_600_, an optical density at 600 nm (OD_600_) of 8.0. The lysed cell suspension was extracted with chloroform-methanol. Briefly, a five-fold volume of chloroform-methanol (2:1; v/v) was added, mixed vigorously for 2 min, and left at room temperature for 10 min. Following the addition of a three-fold volume of chloroform-2% NaCl (1:1; v/v) and centrifugation, the lower layer was recovered and concentrated under vacuum. The lipids were dissolved in chloroform-methanol (2:1; v/v), applied to silica thin-layer chromatography (TLC). The plate was developed with chloroform/methanol/acetic acid (65:25:10, v/v/v) and visualized with CuSO_4_ (100 mg/ml) containing 8% phosphoric acid and heat at 180°C.

### Lipid binding assay

Membrane lipid strips were purchased from Echelon and were pre-spotted with 100 pmol of 15 different lipid species found in cell membranes. To test protein interactions with bacterial membrane lipids, membranes were blocked with TBST containing 1% BSA (w/v) for 1 h at RT. After washing with TBST, membranes were incubated with the MBP-His_6_ or MBP-His_6_-SaeS (2.5 μM each) in TBST containing 1% BSA for 1 h at RT and washed with TBST. The bound protein was detected with a primary α-His_6_ (ThermoFisher) followed by secondary StarBright Blue 700 Goat Anti-Mouse (Bio-Rad). Membranes were visualized using an Amersham ImageQuant800. All Western blots were repeated three times with similar results.

### Quantitative reverse transcription-polymerase chain reaction

To measure mRNA abundance, cells were grown in TSB for 6 hr and total bacterial RNA was isolated using the RNeasy mini kit (Qiagen) with on-column DNA digestion according to the manufacturer’s instructions. After purification, contaminating DNA was removed with RNase-free DNase I. RNA was then purified again using RNeasy Mini columns. cDNA was synthesized with a High Capacity cDNA Reverse Transcription kit (Applied Biosystems) according to the manufacturer’s instructions. Quantification of transcripts was carried out by quantitative reverse transcription polymerase chain reaction (RT-PCR) using SYBR Green PCR Master Mix (Applied Biosystems) in a QuantStudio 6 Flex Real-Time instrument (Applied Biosystems). mRNA abundance was determined by using a standard curve obtained from PCR products generated with serially diluted genomic DNA, and results were normalized to the levels of the *gryB* gene as a reference gene since the abundance of *gyrB* was validated to remain constant (< 0.5 fold change) across all strains and conditions analyzed. Data shown are an average from at least three independent experiments with statistical analysis using Prism 8. Primers used in quantitative RT-PCR assay are presented in table 2.

### Cytotoxicity assay

Bacteria were subcultured for 5 h and 8 h, OD600 normalized, and pelleted by centrifuging at 4,000 rpm for 10 min. The supernatant was filtered through a 0.22-μm filter. Primary human PMNs in RPMI without phenol red (Gibco) supplemented with 10% fetal bovine serum (FBS) were incubated with 2.5% culture filtrate at 37°C and 5% CO_2_. After 1 h, 10 μL of CellTiter 96 AQueous One solution (Promega) was added to each well, and the plate was incubated for another 1.5 h at 37°C and 5% CO2. The absorbance at 490 nm was measured using a PerkinElmer 2103 EnVision multilabel plate reader.

### Animal experiment

For the murine blood infection experiment, overnight cultures of *S. aureus* were inoculated 1:100 into fresh TSB and grown for 2 h with shaking at 37°C. Cells were collected, washed, and suspended in sterile PBS to obtain an inoculum of 1 × 10^9^ CFU mL^−1^. The equal number of male and female of eight-week-old C57BL/6 mice (n=16, 8-week-old, Jackson Laboratory) was infected via retro-orbital injection with 2.5 × 10^7^ CFU of either wild-type, *cls1, cls2*, and *cls1 cls2* mutant strains. Mouse survival was monitored for 22 d. Survival data were analyzed by the log rank (Mantel-Cox) test.

## Acknowledgments

This work was supported by the NIH grant funding AI143792 to TB, AI137403 to SRB and AI137336, AI099394, AI105129 to VJT. The funders had no role in study design, data collection, and interpretation or the decision to submit this work for publication.

